# SARS-CoV-2 Viral Replication in a High Throughput Human Primary Epithelial Airway Organ Model

**DOI:** 10.1101/2021.06.15.448611

**Authors:** Christine R. Fisher, Felix Mba Medie, Rebeccah J. Luu, Robert Gaibler, Caitlin R. Miller, Thomas J. Mulhern, Vidhya Vijayakumar, Elizabeth Marr, Jehan Alladina, Benjamin Medoff, Jeffrey T. Borenstein, Ashley L. Gard

## Abstract

COVID-19 emerged as a worldwide pandemic early in 2020, and at this writing has caused over 170 million cases and 3.7 million deaths worldwide, and almost 600,000 deaths in the United States. The rapid development of several safe and highly efficacious vaccines stands as one of the most extraordinary achievements in modern medicine, but the identification and administration of efficacious therapeutics to treat patients suffering from COVID-19 has been far less successful. A major factor limiting progress in the development of effective treatments has been a lack of suitable preclinical models for the disease, currently reliant upon various animal models and *in vitro* culture of immortalized cell lines. Here we report the first successful demonstration of SARS-CoV-2 infection and viral replication in a human primary cell-based organ-on-chip, leveraging a recently developed tissue culture platform known as PREDICT96. This successful demonstration of SARS-CoV-2 infection in human primary airway epithelial cells derived from a living donor represents a powerful new pathway for disease modeling and an avenue for screening therapeutic candidates in a high throughput platform.

## Introduction

COVID-19 emerged as a worldwide pandemic early in 2020^1^, and has since caused almost 600,000 deaths in the United States and nearly 4 million deaths world-wide^2^. The emergence of multiple safe and efficacious vaccines^3^ has begun to turn the tide of infection and death in regions where the vaccine is widely available. The successful development of these vaccines in a matter of months represents an extraordinary achievement. However, the identification of effective treatments for COVID-19, either from re-purposed drugs or new therapeutic candidates, has lagged severely. A significant factor limiting progress is the absence of preclinical models that are predictive of human disease. Significant progress in the generation of animal models for COVID-19 research has recently been reported^4,5^, but the availability and applicability of these models for wide use in pharmaceutical development remains a challenge. Preclinical cell culture models based on immortalized cell lines such as Vero E6 and A549 have also been reported^6,7^, but lack key aspects of the host biology such as ciliary differentiation and mucociliary clearance believed to be critical in understanding pathogenesis and predicting therapeutic efficacy. Human primary airway epithelial cell culture at an air-liquid interface (ALI) represents a powerful platform technology for respiratory research^8^, with applications ranging from chronic pulmonary conditions to acute respiratory infections and toxic exposures to the lung. Recently, a Transwell™-based assay comprising human primary airway epithelial cell culture has also been used to investigate SARS-CoV-2 infection^9^. However, such membrane insert-based systems are quite limited in several key aspects, such as difficulties in high resolution imaging, large media volume, and the absence of dynamic media circulation and real-time sensing, relative to the organ-on-chip platform presented here.

In response to these challenges, the field of organs-on-chips has turned intense attention toward modeling of respiratory pathogens such as influenza and SARS-CoV-2, and a number of groups have reported results relevant to the effort to establish therapeutic screening tools for these diseases^10–13^. These platforms utilize human primary alveolar or airway epithelial cells in microfluidic organ models designed to closely mimic not only the relevant cell biology but other key aspects of the *in vivo* microenvironment. To date, these studies have been limited to analysis of SARS-CoV-2 pseudoviruses due to significant challenges in both accessing and executing organ-on-chip experiments in BioSafety Level 3 (BSL-3) facilities. While pseudoviruses represent a potentially useful and convenient model for the study of viral entry, several key studies of neutralizing antibodies against SARS-CoV-2^14^ and in particularly highly transmissible variants^15^ indicate that they do not always accurately reflect neutralization of the live virus. Furthermore, in lacking the viral life cycle of wildtype viruses, pseudoviruses are less suitable screening tools for compounds that may target stages other than viral entry. This and other related considerations spurred our efforts to investigate viral infection kinetics of the live SARS-CoV-2 virus in the PREDICT96-ALI platform in a BSL-3 laboratory environment.

Here we report the first demonstration of SARS-CoV-2 infection and viral replication in a human primary cell-based airway-on-a-chip organ model. We utilized the PREDICT96-ALI^13^ system that has previously been demonstrated as a platform for the study of infection human airway tissue models with influenza virus and hCoV-NL63, in collaboration with the University of Massachusetts Medical School under the DARPA PREPARE program. Unlike earlier studies, this work investigates the inoculation of live SARS-CoV-2 virus into human primary airway epithelial cells in a BSL-3 environment, an important step forward in the arsenal of preclinical tools for modeling COVID-19 infection. This report is based on an initial set of experiments with a single donor population of human primary airway epithelial cells. Further studies with additional donor cells are underway, but this first successful demonstration of SARS-CoV-2 infection in a human primary organ-on-chip model represents an exciting milestone and opportunity for modeling pandemic respiratory infections. The streamlined applicability of this platform for BSL-3 operation, combined with its inherent high throughput nature, integrated perfusion, *in situ* imaging, and sensing capabilities^16,17^ render PREDICT96 a powerful tool for modeling respiratory infections, and an avenue for screening therapeutic candidates for treatment of these severe diseases in a rapid and efficient manner.

## Materials and Methods

### Microfluidic Platform and Integrated Micropumps

The PREDICT96 organ-on-chip platform consists of a microfluidic culture plate with 96 individual devices and a perfusion system driven by 192 microfluidic pumps integrated into the plate lid^17^. As described in our previous work^13,16–18^, in order to establish recirculating flow in the microchannels, we incorporate a self-priming micropump array into the lid that serves as the fluidic interface with the culture plate via stainless steel tubes. The PREDICT96 pumping system has 192 individual pneumatically-actuated micropumps embedded in the plate lid: two per culture device, one for the chamber above the membrane and one for below. However, since the upper chamber is at air-liquid interface (ALI), only the 96 pumps serving the microchannel are in use during experiments. Actuation of the pumps transfers media between the wells linked by the bottom microfluidic channel and establishes a hydrostatic pressure differential, inducing flow through each microchannel.

### Collection and Preparation of Human Primary Bronchial Epithelial Cells from Healthy Living Donors

Human small airway epithelial cells were isolated from living donors by research bronchoscopy at Massachusetts General Hospital (MGH), Boston, MA. Bronchoscopy brushings were collected into collection media (RMPI (ThermoFisher) + 2% human serum albumin (ThermoFisher) + 5 μM ROCKi (Tocris)) and transported on ice to Draper (Cambridge, MA)^19^. All research involving human subjects was approved by the MGH and Partners Healthcare System Institutional Review Board, and adhered to all applicable institutional and sponsor ethical guidelines, including but not limited to anonymization of donors for personal identity. All methods were in adherence with applicable safety and laboratory practice guidelines.

### Culture, Cryopreservation, and PREDICT96-ALI Seeding and Differentiation of NHBEs

After isolation and transport to Draper, human small airway epithelial cells were spun down at 200 x *g* for 5 min, re-suspended in SAGM + 4i media, and transferred to a well plate coated with 804G media (804G media production described previously^19,20^). Donor cells reported in this manuscript were derived from donor DH01: 28 years, female, Asian (non-Hispanic). Cells were cultured in SAGM + 4i media^13^, and passaged using Accutase (Sigma). After 4 passages, cells were cryopreserved in Cryostor 10 (STEMCELL Technologies). PREDICT96-ALI plates were prepared as previously described^13^. DH01 cells were thawed and plated at 15,000 cells per device in a 3 μL volume directly onto the membrane. Cell/tissue growth in submerged and then ALI conditions was conducted as previously reported in a custom-ALI media^13^. Tissues were matured over the course of 4 weeks in ALI culture prior to viral infection experiments. To remove accumulated mucus from the apical surface of the maturing tissue, 100 μL of 1x Hank’s Balanced Salt Solution (HBSS, Sigma) was added to the apical surface of each tissue and incubated on the tissues for 1 h at 37 °C with rocking, subsequently followed by an additional 5 min wash with 100 μL of 1x HBSS at room temperature every 7 days (d). Downstream processing of tissues following viral inoculation included tissue apical washes, basal media collection, harvest of tissue for RNA extraction, and fixation for immunofluorescence imaging.

### Immunofluorescence and Confocal Imaging

Immunofluorescence staining was conducted as previously reported^13^. Primary antibodies were used to stain basal cells (CK5, Thermo Fisher), goblet cells (Muc5A, Thermo Fisher), ciliated cells (acetylated-tubulin, Abcam) and SARS-CoV-2 (Spike and Nucleocapsid proteins, GeneTex). The SARS-CoV-2 Spike and Nucleocapsid protein antibodies were combined when added to the PREDICT96-ALI airway tissues during primary antibody incubation steps. Secondary antibodies used were Alexa Fluor 488 goat anti-mouse IgG (Thermo Fisher) and Alexa Fluor 555 goat anti-rabbit IgG (Thermo Fisher). Tissues were incubated with counter-stains Hoechst 33342 dsDNA nuclear stain (ThermoFisher) and Phalloidin-iFluor 647 reagent (Abcam). Immunofluorescent labeled PREDICT96-ALI tissues were imaged *in situ* within the PREDICT96-ALI plate by way of the optical qualities of the PREDICT96-ALI platform and using a Zeiss LSM700 laser scanning confocal microscope and Zen Black software (Zeiss). Tile scans and z-stacks of the tissues at 40x and 10x magnification were acquired.

### Mean Fluorescence Intensity Quantification

10x tile scans of the middle of each PREDICT96-ALI tissue device were captured on a Zeiss LSM700 laser scanning confocal microscope with Zen Black software (Zeiss). The MFI of the signal from the red channel, or the Nucleocapsid and Spike protein expression, of these regions was measured via ImageJ Fiji. The average signal from the secondary only controls were subtracted to remove the background signal, and the average MFI from each infection condition was calculated.

### Club-Cell Secretory Protein (CCSP) Luminex

After 4 weeks of ALI culture of DH01 human primary airway epithelial cells, basal media samples were collected for quantification of soluble CCSP via a premade Magnetic Luminex Performance Assay kit (Uteroglobin/SCGB1A1, R&D Systems). Culture media was collected from the bottom chamber of each tissue device and immediately frozen at −20°C until use. Undiluted media samples were processed according to the manufacturer’s protocol and analyzed via a Luminex FLEXMAP 3D. The data collected were used to generate standard curves for each analyte using a five parameter logistic (5-PL) curve-fit to determine the concentration of CCSP in the sample. Concentration of CCSP concentration was measured for each individual PREDICT96-ALI microtissue device, and devices derived from the same experimental conditions were averaged together and reported as mean ± standard deviation (N=6 devices).

### Inoculation of PREDICT96-ALI Tissues with SARS-CoV-2

SARS-CoV-2 strain USA-WA1/2020, passage 4 virus was obtained from Joseph Brewoo and Sam R. Telford III (New England Regional Biosafety Laboratory, Global Heath & Infectious Disease, Tufts University). All of the work with SARS-CoV-2 was performed at BSL-3. Inoculation of PREDICT96-ALI airway tissues with SARS-CoV-2 (and mock control) was performed similarly to coronavirus infections described previously^13^. Briefly, the PREDICT96-ALI tissues were prepared by performing a mucous wash with Hanks’ Balanced Salt Solution (HBSS). Freshly-thawed SARS-CoV-2 was diluted in infection media (DMEM (Gibco) supplemented with 0.1% bovine serum albumin (Sigma), 1x Penicillin/Streptomycin (Gibco), 1x non-essential amino acids, NEAA(Gibco)) to MOI 0 (mock control), 0.025 or 0.0025. Viral inoculum (or mock control) was incubated on the apical surface of the microtissues for 2 h rocking at 34°C. Inoculum was subsequently removed, and both the top and bottom chambers of each microtissue device were washed 4x with HBSS. The final wash reserved as a 0 days post infection (d p.i.) sample. Fresh custom-ALI media was introduced into the basal ports before static incubation at 34°C.

On 2, 4, and 6 d p.i., the apical surface of the tissues and bottom chamber of each tissue were washed twice with HBSS (40 min per wash). The two HBSS washes collected from the apical surface of a single device were pooled and stored at −80°C until downstream assessment. Fresh custom-ALI media was introduced into the bottom chamber before returning the plate to 34°C. At the conclusion of the experiment on day 6 p.i., the PREDICT96-ALI airway tissues were fixed with 4% paraformaldehyde (Electron Microscopy Sciences) for 1 h at room temperature. Tissues were then washed with and incubated in phosphate buffered saline (PBS) until immunofluorescence staining.

### RNA extraction and RT-qPCR

Viral RNA was extracted from HBSS wash samples and tissue RNA was extracted from microtissue devices as previously described^13^. Briefly, QIAamp Viral RNA Mini Kits (Qiagen) and RNeasy Micro Kits (Qiagen) were used for RNA extraction following the manufacturer’s protocols. Uniform volumes of HBSS wash samples were extracted and eluted in 60 μl AVE buffer, and individual PREDICT96-ALI microtissues were extracted in RLT buffer with 0.01% v/v 2-mercaptoethanol (Sigma) and eluted in 14 ul RNAse-free water. One-step reverse transcription quantitative polymerase chain reaction (RT-qPCR) was then performed on extracted RNA samples using a QuantiTect Probe RT-PCR kit (Qiagen) with 7.8 μl purified viral RNA or specified quantities (2.5 ng or 100 ng) of tissue RNA in a 20 μl reaction. The reaction was run in an Applied Biosystems QuantStudio 7 Flex System (Thermo Scientific) using the following condition: 50°C for 30 min, 95°C for 15 min, 45 cycles of 94°C for 15 sec and 60°C for 60 sec. The following primers and probes targeting SARS-CoV-2 N protein (N1 and N2) were used to detect viral RNA in HBSS wash samples and bulk tissue RNA: nCOV_N1 Forward Primer Aliquot, 100 nmol (IDT, 10006830), nCOV_N1 Reverse Primer Aliquot, 100 nmol (IDT, 10006831), nCOV_N1 Probe Aliquot, 50 nmol (IDT, 10006832), nCOV_N2 Forward Primer Aliquot, 100 nmol (IDT, 10006833), nCOV_N2 Reverse Primer Aliquot, 100 nmol (IDT, 10006834), and nCOV_N2 Probe Aliquot, 50 nmol (IDT, 10006835). Additional primers and probe targeting human RNase P were used as an internal control to verify that nucleic acid is present (positive extraction control) for assessment of bulk tissue RNA: RNase P Forward Primer Aliquot, 100 nmol (IDT, 10006836), RNase P Reverse Primer Aliquot, 100 nmol (IDT, 10006837), and RNase P Probe Aliquot, 50 nmol (IDT, 10006838). TaqMan Gene Expression Assays (Thermo Scientific) were used to detect mRNA transcript targets in microtissues: ACE2 (Hs01085333_m1), TMPRSS2 (Hs01122322_m1), SCGB1A1(Hs00171092_m1), Muc5AC (Hs01365616_m1), FoxJ1 (Hs00230964_m1), TP63 (Hs00978340_m1). Absolute quantification (copies/mL) of SARS-CoV-2 N1 and/or N2, as well as RNase P control) of wash samples and tissue RNA were calculated using a standard curve generated from serial dilutions of linearized SARS-CoV-2 viral RNA (American Type Culture Collection, ATCC). Quantification of microtissue transcript expression were calculated using the Comparative cycle threshold (Ct) values were determined using the method described by Schmittgen and Livak^21^ using normalization to the housekeeping gene GAPDH.

Washes from 6x PREDICT96-ALI devices per condition per time-point were tested by RT-qPCR in duplicate. Samples with cycle threshold (ct) values above 37 were omitted. Absolute quantification (copies/mL) of supernatant viral RNA was calculated using a standard curve generated from serial dilutions of 2019-nCoV_N_Positive Control viral RNA (IDT, 10006625).

### Plaque assay

Wash samples collected from inoculated PREDICT96-ALI plates were titered via plaque assay as previously reported^22^. Briefly, Vero E6 cells (ATCC) were plated in 6-well plates and incubated at 37°C and 5% CO2 until confluent. Viral samples were diluted in DMEM with 0.1% BSA, 1x NEAA and 1x Penicillin/Streptomycin: 1:10 for mucous wash samples from the infected PREDICT96-ALI tissues and ten-fold serial dilutions for stock virus. Wells were washed with PBS then incubated with viral dilutions (300 μl total) for 2 h at 37°C, rocking every 15 min. Plates were then washed with PBS, overlaid with DMEM supplemented with final concentrations of 2% fetal bovine serum (FBS), 1x NEAA, 1x Penicillin/Streptomycin, and 1.2% Cellulose (colloidal, microcrystalline (Sigma)), and incubated for 3 days. After removal of the overlay, plates were washed with PBS, fixed with 4% PFA for 30 min, stained with a 0.1% crystal violet (Sigma)/4% ethanol solution, and imaged. Plaque forming units/milliliter (PFU/mL) were calculated as follows: number of plaques / (dilution x total inoculum volume (mL)).

### Statistical analysis

Data are presented as mean ± standard deviation or standard error of the mean (indicated in figure legends) and were analyzed using Graphpad Prism version 9.0.0 for Windows (GraphPad Software). Statistical significance was determined using one- or two-way analysis of variance (ANOVA) with Tukey’s post hoc test for multiple comparisons, where appropriate. A p-value lower than 0.05 was considered statistically significant.

## Results

### PREDICT96-ALI supports formation of mature human airway tissue from research bronchoscopy-derived donor cells

Human tissues differentiated and maintained in PREDICT96-ALI were derived from freshly harvested human bronchial epithelial cells from living research bronchoscopies, as previously reported^13^. Herein, all human tissues in PREDICT96-ALI were derived from donor DH01: healthy, 28 year old, female, Asian non-Hispanic. The airway tissue used in the present SARS-CoV-2 infection studies resembled previously reported^13^ pseudostratified, 30-50 μm thick tissues matured over 4-weeks in the PREDICT96-ALI platform. Immunofluorescent staining confirmed the relative populations of ciliated cells (acetylated-tubulin), basal cells (CK5), and goblet cells (Muc5AC) within the mature tissues, all with scale bar = 50 μm **(Figure 1A)**. Similarly, tissue function was monitored and club cell presence in the tissues determined by measuring CCSP secretion into the media in the bottom chamber of each device via Luminex at 4 weeks of ALI **(Figure 1B)**.

**Figure 1:**
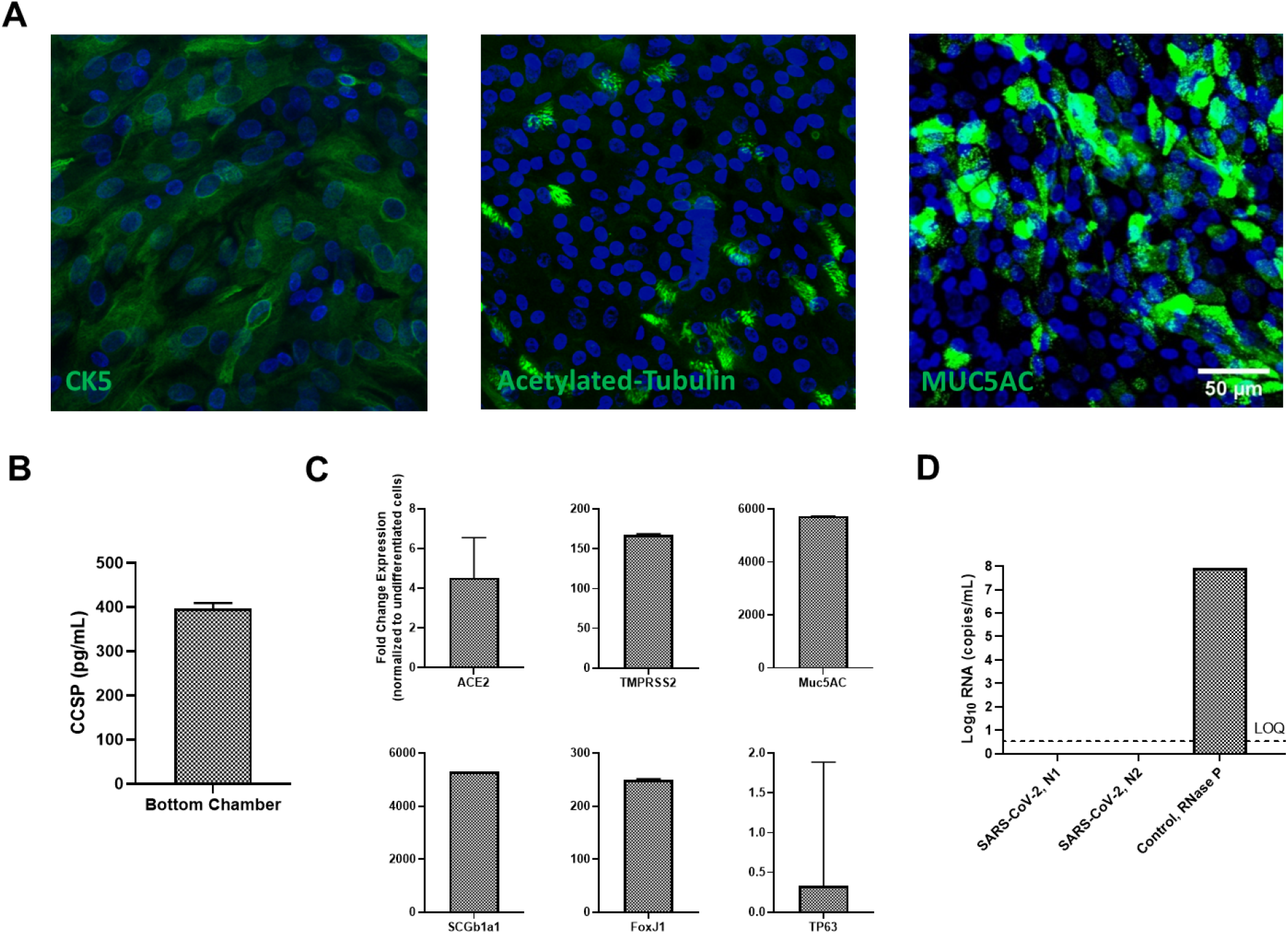
Donor cells derived from research bronchoscopy and differentiated in PREDICT96-ALI exhibit structure and composition of mature human airway. **(A)** Representative immunofluorescence images demonstrating the presence of basal (CK5, green), ciliated (acetylated-tubulin, green), and goblet (Muc5AC, green) cells in healthy, uninfected pseudostratified PREDICT96-ALI airway tissue derived from donor DH01 and grown to 4-weeks ALI, shown with a 50 μm scale bar. Images are counter-stained for dsDNA nucleic acids (Hoechst 33342, blue) **(B)** Club Cell Secretory Protein (CCSP) detection in PREDICT-ALI airway experiments containing donor DH01 at 4 weeks ALI. Media samples collected from the bottom microfluidic chamber of each PREDICT96-ALI tissue exhibit detectable levels of CCSP (395.88 ± 13.625 pg/mL), indicating a robust population of functioning secretory club cells. N = 4 tissue media samples selected at random from two independent, representative PREDICT96-ALI experiments. Data presented as mean concentration (pg/mL) and error bar represents standard error of the mean. **(C)** Relative fold change in expression of key mRNA transcripts from differentiated airway tissues in PREDICT96-ALI derived from donor DH01. Reverse transcription quantitative polymerase chain reaction (RT-qPCR) was used to detect transcripts for ACE2 and TMPRSS2, both critical for establishing SARS-CoV-2 infection. Transcripts were also detected for proteins associated with mature airway epithelial cell populations: Muc5AC, SCGB1A1, TP63 and FoxJ1 for goblet, club, basal and ciliated cells, respectively. Comparative Ct values were determined using the method described by Schmittgen and Livak^21^, used to calculate the relative quantification of gene expression using GAPDH as a reference gene, and normalized to undifferentiated DH01 basal epithelial cells. N = 12 healthy tissue replicates selected at random from one representative PREDICT96-ALI experiment. Data presented as fold change expression and error bar represents standard error of the mean. **(D)** Diagnostic confirmation that isolated epithelial cells from donor DH01 are SARS-CoV-2 negative. Negativity was determined by RT-qPCR using primer and probe sets for the detection of two regions in the SARS-CoV-2 nucleocapsid (N) gene (N1 and N2) and one primer and probe set to detect human RNase P. Limit of quantification (LOQ) line indicates copy number corresponding to Ct values of ≥ 40 cycles. Samples that did not meet the minimum signal intensity (undetermined) after 40 PCR cycles are excluded. Data presented as Log10 viral RNA copies/mL and error bar represents standard error of the mean.

In preparation for challenging the PREDICT96-ALI airway tissues with SARS-CoV-2 virus, the relative mRNA transcript expression of angiotensin-converting enzyme 2 (ACE2), which is the receptor used by SARS-CoV-2, and transmembrane protease serine 2 (TMPRSS2), which primes the spike protein of SARS-CoV-2, were established in the healthy, mature uninfected tissues. In addition, key gene transcripts associated with differentiated airway were confirmed and normalized to GAPDH and undifferentiated basal cells: Muc5AC (goblet cells), SCGB1A1 (club cells), FoxJ1 (ciliated cells), and TP63 (basal cells) **(Figure 1C)**. Moreover, the airway tissues displayed robust RNase P transcript quantities and had no measurable levels of SARS-CoV-2 N1 and N2 viral proteins detected prior to inoculation with wild-type SARS-CoV-2 **(Figure 1D)**. Altogether, the PREDICT96-ALI tissues were in a mature, differentiated state and negative for SARS-CoV-2 prior to initiation of SARS-CoV-2 infection studies at 4 weeks of ALI.

### PREDICT96-ALI airway tissue supports inoculation with wildtype SARS-CoV-2 virus and yields replicative virus

Next, we examined the effect of inoculation of mature PREDICT96-ALI airway tissue with wildtype SARS-CoV-2 USA-WA1/2020. RNA specific to SARS-CoV-2 N1 was detected in tissues in both infected conditions (MOI 0.025 and 0.0025) relative to mock control, 0 MOI **(Figure 2A)**. While RNA levels tapered off in the lower MOI condition and were undetectable by day 6 post infection (p.i.), RNA copy numbers increased 600-fold between days 0 and 6 in apical tissue washes from the 0.025 MOI infection condition. Viral copy numbers were comparable to prior reports in other *in vitro* primary airway models and *in vivo* clinical specimens (sputum, swab) from patients with active virus replication in tissues of the upper respiratory tract^23,24^. Similar trends were observed when the RNA was assayed with SARS-CoV-2 N2-specific probes (data not shown).

**Figure 2:**
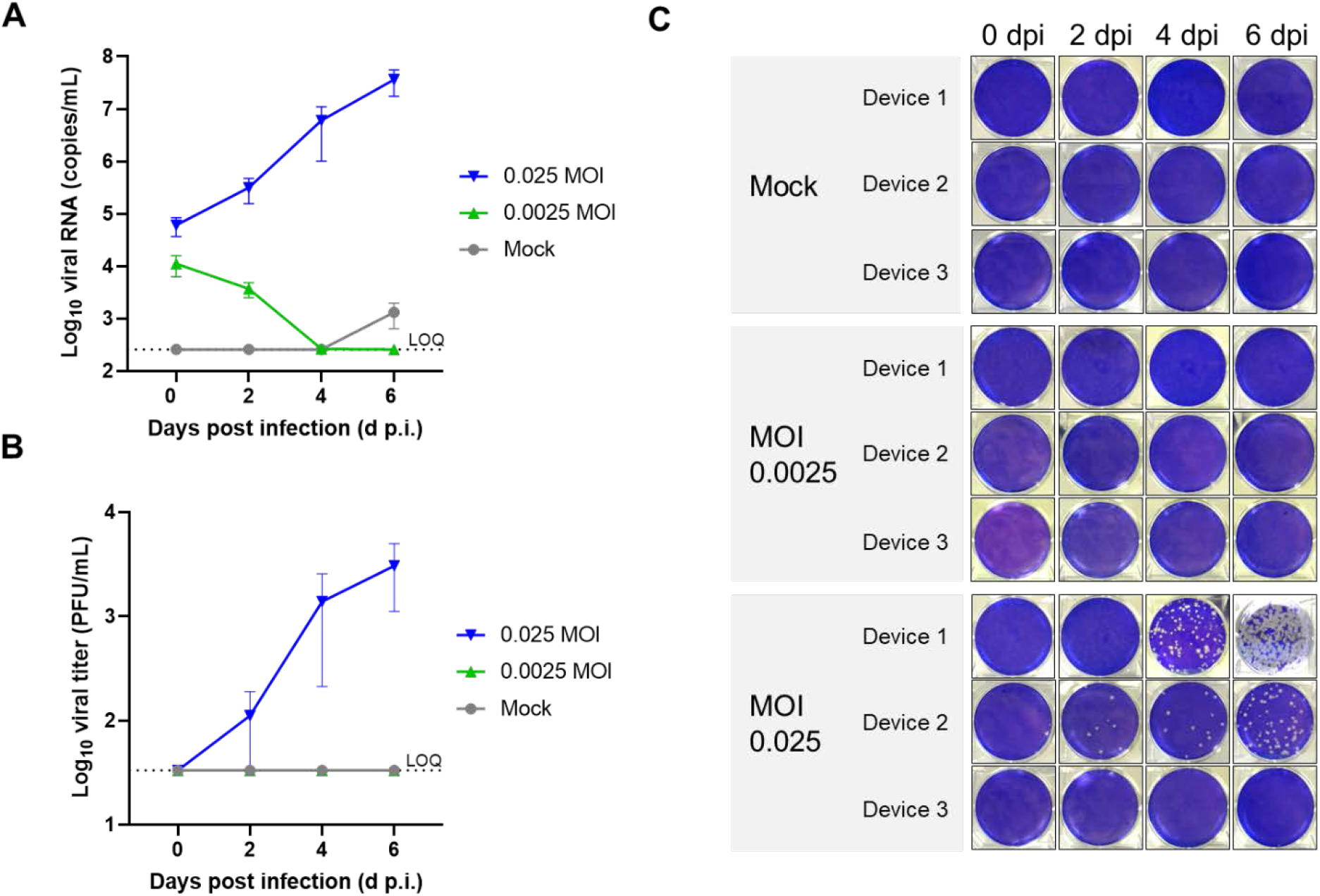
SARS-CoV-2-inoculated PREDICT96-ALI airway tissues yield replicative virus by RT-qPCR and plaque assay. **(A)** RT-qPCR analyses for viral RNA copies in the apical wash of PREDICT96-ALI airway tissue at 0, 2, 4, and 6 days post infection (d p.i.). N = 6 tissue replicates per time-point and condition from one independent experiment; symbols represent mean and error bars represent standard error of the mean. Limit of quantification (LOQ, dotted line) indicates copy number corresponding to Ct values ≥ 37 cycles. Replicates that did not meet the minimum signal intensity after 37 PCR cycles are displayed along the LOQ line. **(B)** Plaque assay analysis detecting replicative virus from apical wash samples of PREDICT96-ALI airway tissue at 0, 2, 4, and 6 d p.i. N = 3 tissue replicates per time-point and condition from one independent PREDICT96-ALI experiment. Plaque forming units (PFU) were calculated and quantified from plaque assay plate images. Data is presented as Log10 viral titer PFU/mL and error bars represent standard error of the mean. Limit of quantification (LOQ, dotted line): ≤ 33.3 PFU/mL. Replicates that did not generate plaques are displayed along the LOQ line. **(C)** Macroscopic images of 6-well plaque assay wells corresponding to data displayed in panel B.

Apical tissue wash samples taken throughout the infection study were also assessed for replication-competent virus by standard plaque assay **(Figure 2B-C)**. Plaques were detectable as early as day 2 p.i. in the 0.025 MOI-infected condition. While there was substantial variability in titer among the three 0.025 MOI-infected devices tested, the overall trend matched that of the RT-qPCR data and prior reports in other *in vitro* primary airway models^9,25^, further validating the model’s capacity to support SARS-CoV-2 replication.

### PREDICT96-ALI airway tissue exhibits heterogeneous SARS-CoV-2 viral foci

SARS-CoV-2 infection in PREDICT96-ALI airway tissues was additionally characterized by immunofluorescence microscopy. **Figure 3A** shows max projection images of immunofluorescence (IF) staining of PREDICT96-ALI tissues devices inoculated with various MOIs of SARS-CoV-2. The images exhibit greater abundance of SARS-CoV-2 spike and nucleocapsid proteins (red) as MOI increases. Strong expression of nucelocapsid and spike proteins is seen at 6 days p.i. and distributed within the 0.025 MOI tissues. Foci range in size from single cells to large multi-cell clusters indicating SARS-CoV-2 replication and spread to neighboring cells, an observation also reported in prior studies working with *in vitro* primary upper airway models^25^. MFI quantification of SARS-CoV-2 infection shows significant expression of SARS-CoV-2 positive cells at 0.025 MOI **(Figure 3B)**. MFI quantification also detected inter-device variation in SARS-CoV-2 positivity at 6 d p.i, similar to the variability exhibited between devices in the plaque assay **(Figure 2C)**. While the relatively low MOI used likely influenced the variability, its biological properties and mechanisms are unknown and actively being investigated.

**Figure 3:**
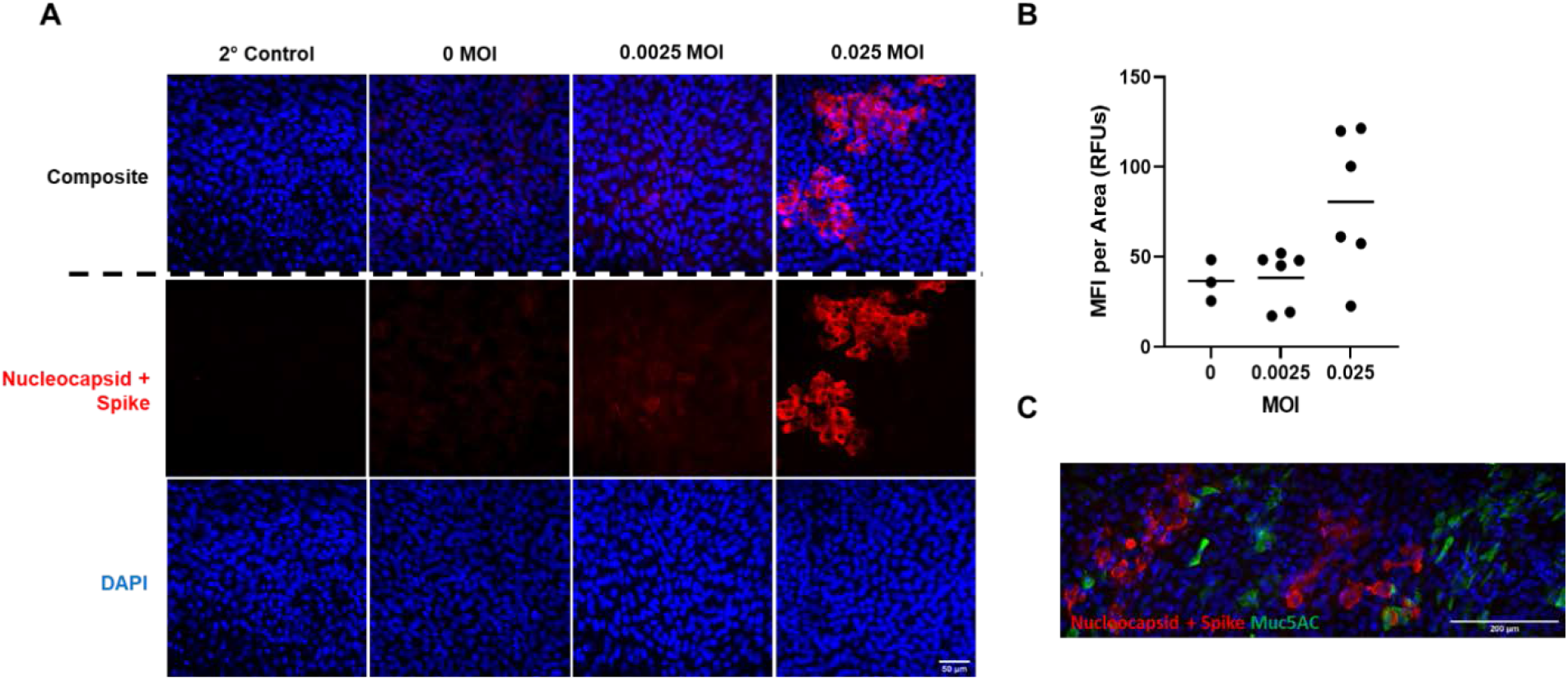
Immunofluorescent staining of SARS-CoV-2-inoculated PREDICT96-ALI airway tissues exhibits discreet SARS-CoV-2 viral foci. **(A)** Max projection immunofluorescent images depicting positive staining of the SARS-CoV-2 spike and nucleocapsid proteins (S and N, respectively, red) and dsDNA nucleic acids (Hoechst 33342, blue) within PREDICT96-ALI airway tissue from donor DH01. Tissues were fixed with 4% paraformaldehyde at 6 d p.i. following inoculation with SARS-CoV-2 USA-WA1/2020 (MOI 0, 0.0025 or 0.025). Secondary antibody control showed low background noise. Images were taken at 40x magnification and are shown with a 50 μm scale bar. **(B)** Mean Fluorescent Intensity (MFI) image quantification of SARS-CoV-2 infection in PREDICT96-ALI airway tissues associated with panel A. MFI from the red channel (S and N proteins) indicate a trend of increased signal in the 0.025 MOI condition, but is not significant compared to the 0 or 0.0025 MOI conditions. The average MFI of the 0, 0.0025, and 0.025 MOI are 36.67±11.45, 38.38±15.75, 80.52±39.69 RFUs (mean ± standard deviation), respectively. Images were taken at 10x magnification and analyzed for MFI using ImageJ Fiji. **(C)** Representative max projection immunofluorescent image of SARS-CoV-2 positivity (S and N proteins, red), goblet cells (Muc5AC, green) and dsDNA nucleic acids (Hoechst 33342, blue) within PREDICT96-ALI airway tissue from donor DH01 at 6 d.p.i. following inoculation with SARS-CoV-2 at 0.025 MOI. Image was taken at 40x magnification and is shown with a 200 μm scale bar.

In **Figure 3C**, a max projection 40x tile image depicts SARS-CoV-2-infected PREDICT96-ALI tissue (red) counter-stained with Muc5AC (green) for goblet cells. The image highlights heterogeneous SARS-CoV-2 viral foci and co-localization between SARS-CoV-2 staining to airway goblet cells, indicating that secretory cells are likely one of the target respiratory epithelial cell types of SARS-CoV-2, as reported by other prior studies^23,26^. Cytopathic effect (CPE) is also observed in SARS-CoV-2-infected tissues, as indicated by the irregular morphology observed in some SARS-CoV-2 positive cells **(Figure 3C)**.

Future work will investigate MOIs > 0.025, the cellular tropism of SARS-CoV-2 during the time course of infection, and longer functional experimental infection timelines (> 6 d p.i, an arbitrarily selected terminal time-point) for longitudinal analyses and extended therapeutic evaluation runways. Altogether, these results demonstrate that PREDICT96-ALI is a capable platform for modeling SARS-CoV-2 infection in primary human airway tissue derived from bronchoscopy-derived donor sources and provides the throughput needed to evaluate a wide-range of infection parameters and observe critical insights into the pathogenesis of SARS-CoV-2.

## Discussion

Preclinical animal models and cell culture platforms represent critically important tools in modeling respiratory viruses and predicting efficacy of candidate therapeutics, yet they lack critical features necessary to accurately recapitulate human-specific mechanisms of viral infection and responses to antiviral therapies. Organ-on-chip technologies have the potential to overcome these limitations by offering human-relevant physiological fidelity in infection-based assays. However, these platforms have historically been challenged by excessive complexity and difficulty in direct application in pharmaceutical laboratory environments. Further, in the case of SARS-CoV-2 research, these models have so far been limited to the study of pseudovirus entry into cells rather than direct investigations of infection by the live virus. Here, we report the first SARS-CoV-2 infection study in a human primary cell-based airway-on-a-chip model. Further, we utilize human airway epithelial cells isolated from living donors via research bronchoscopies, a capability that provides the opportunity for evaluating mechanisms of infection for specific patient subpopulations. Our high throughput PREDICT96-ALI platform was successfully adapted to operation in a BSL-3 environment while retaining key biological attributes of functional respiratory tissue. Importantly, PREDICT96-ALI has the demonstrated capacity for use in high-throughput assays, a critical capability for practical drug discovery and therapeutic screening applications.

Further development of this platform technology can enable the study of respiratory infections across large numbers of independent human cell donors, spanning a wide-range of demographics and co-morbidities in order to capture natural variability in the population. Leveraging all of its features, the PREDICT96-ALI model will facilitate high-throughput, preclinical evaluation of therapeutic candidates in a dynamic tissue microenvironment that is receptive to real-time, non-invasive sensing and imaging. These capabilities will be critical to achieving rapid and targeted evaluation of efficacious medical countermeasures.

## Author Interests

All authors declare no conflict of interests.

## Author Contributions

Project design was done by ALG, CRF, and RJL. Design and execution of the experiments was led by ALG along with CRF, FMM, RJL, RG, CRM, TJM, VV, EM, JA, and BM. Data analysis was done by ALG, CRF, FMM, RJL, RG, CRM, TJM, and JTB. The manuscript was written by ALG, CRF, JTB, RJL, RG, and FMM, and reviewed and edited by all authors.

## Acknowledgements

The authors gratefully acknowledge Roger Odegard, Joseph Charest, Rob Larsen, Rachel Fezzie, Amy Duwel and Alla Gimbel, for their technical and programmatic support for this work, Rick Crocker for his support of the laboratory infrastructure, John Gilmartin from the Draper Safety Office, and Tim Petrie for key technical guidance and critical review of the manuscript. We gratefully acknowledge the technical and programmatic support of Sam Telford, Joseph Brewoo and the staff at Tufts NERBL. Funding from Internal Draper Independent Research and Development (IR&D) is gratefully acknowledged.

